# AEGIS: An In Silico Tool to model Genome Evolution in Age-Structured Populations

**DOI:** 10.1101/646877

**Authors:** William J. Bradshaw, Arian Šajina, Dario Riccardo Valenzano

## Abstract

AEGIS (Ageing of Evolving Genomes In Silico) is a versatile population-genetics numerical-simulation tool that enables the evolution of life history trajectories under sexual and asexual reproduction and a wide variety of evolutionary constraints. By encoding age-specific survival and reproduction probabilities as discrete genomic elements, AEGIS allows these probabilities to evolve freely and independently over time. Simulation of population evolution with AEGIS demonstrates that ageing-like phenotypes evolve in stable environments under a wide range of conditions, that life history trajectories depend heavily on mutation rates, and that sexual populations are better able to accumulate high levels of beneficial mutations affecting early-life survival and reproduction. AEGIS is free and open-source, and aims to become a standard reference tool in the study of life-history evolution and the evolutionary biology of ageing.

## Introduction

Species in nature vastly differ in life histories, with dramatic variation in maturation rate, lifespan, and fecundity. In general, age-dependent mortality increases as a function of age while age-dependent fecundity declines, a phenomenon known as *ageing* or *senescence*. However, in some organisms mortality decreases or remains constant through life, while fecundity remains constant or increases (Jones et al., 2014). These difference in demography can have important effects on fitness, giving rise to dramatic differences in lifetime reproductive output between species.

The evolution of age-dependent changes in mortality and reproduction has been an important object of theoretical investigation since the dawn of population genetics, giving rise to a number of theories to explain the widespread occurrence of senescence in nature. Work from Haldane, Medawar, Hamilton and others predicts that the declining force of natural selection after reproductive maturation should inevitably lead to the accumulation of deleterious gene variants, resulting in increased mortality later in life (Haldane, 1941; Medawar, 1952; Hamilton, 1966; Charlesworth, 2000). While these mutation-accumuation theories of ageing explain ageing as a fundamentally non-adaptive process, other evolutionary theories of ageing suggest senescence could evolve as an antagonistic side-effect of positively-selected traits (Williams, 1957), or even as a kin- or group-selected adaptation in its own right (Longo et al., 2005; Lohr et al., 2019).

Up to now, the evolution of life-history traits, including age-dependent changes in survival and reproduction, has primarily been performed using analytical approaches (Hamilton, 1966; Charlesworth, 1994; Fisher, 1930); while some simple numerical models exploring the evolution of ageing have been proposed (Penna, 1995; Dzwinel et al., 2005; Werfel et al., 2015), there remains a need for a flexible simulation tool to model the evolution of ageing. In particular, a model which permits independent evolution in both mortality and fecundity at different ages could capture a wider range of possible life histories and so provide a particularly powerful tool for simulating the evolution of ageing.

Here, we present and release AEGIS (Ageing of Evolving Genomes *In Silico*), a ready-to-use numerical model of genome evolution that simulates how age-dependent changes in survival and reproduction evolve under a range of different ecological and demographic scenarios.

## New Approaches

AEGIS is a Python-based platform implementing and extending a discrete-time, non-spatial numerical model of genome evolution (Šajina et al., 2016). In this model, each individual is represented by a diploid bit-string genome, which is divided into age-specific survival and reproduction loci specifying the baseline survival and reproduction probabilities of that individual at the appropriate age (Fig. 1A), where “age” designates the number of discrete-time stages since the individual was added to the population. These probabilities scale linearly between user-specified bounds (*p*_min_, *p*_max_) based on the additive sum *L* of the bit values in the appropriate loci across both chromosomes:

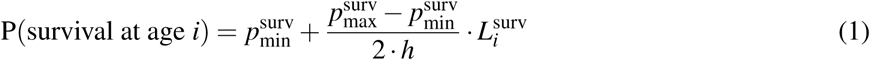

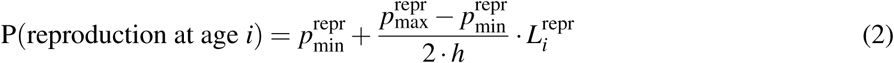

where *h* is the number of bits per locus per chromosome. The survival and reproduction probabilities are therefore lowest when all bits in the corresponding loci are equal to 0, and highest when they are all equal to 1. In addition to survival and reproduction loci, the genome also contains some number of neutral loci without a phenotypic effect, which serve to track the effects of neutral evolution on genome composition.

**Figure 1:**
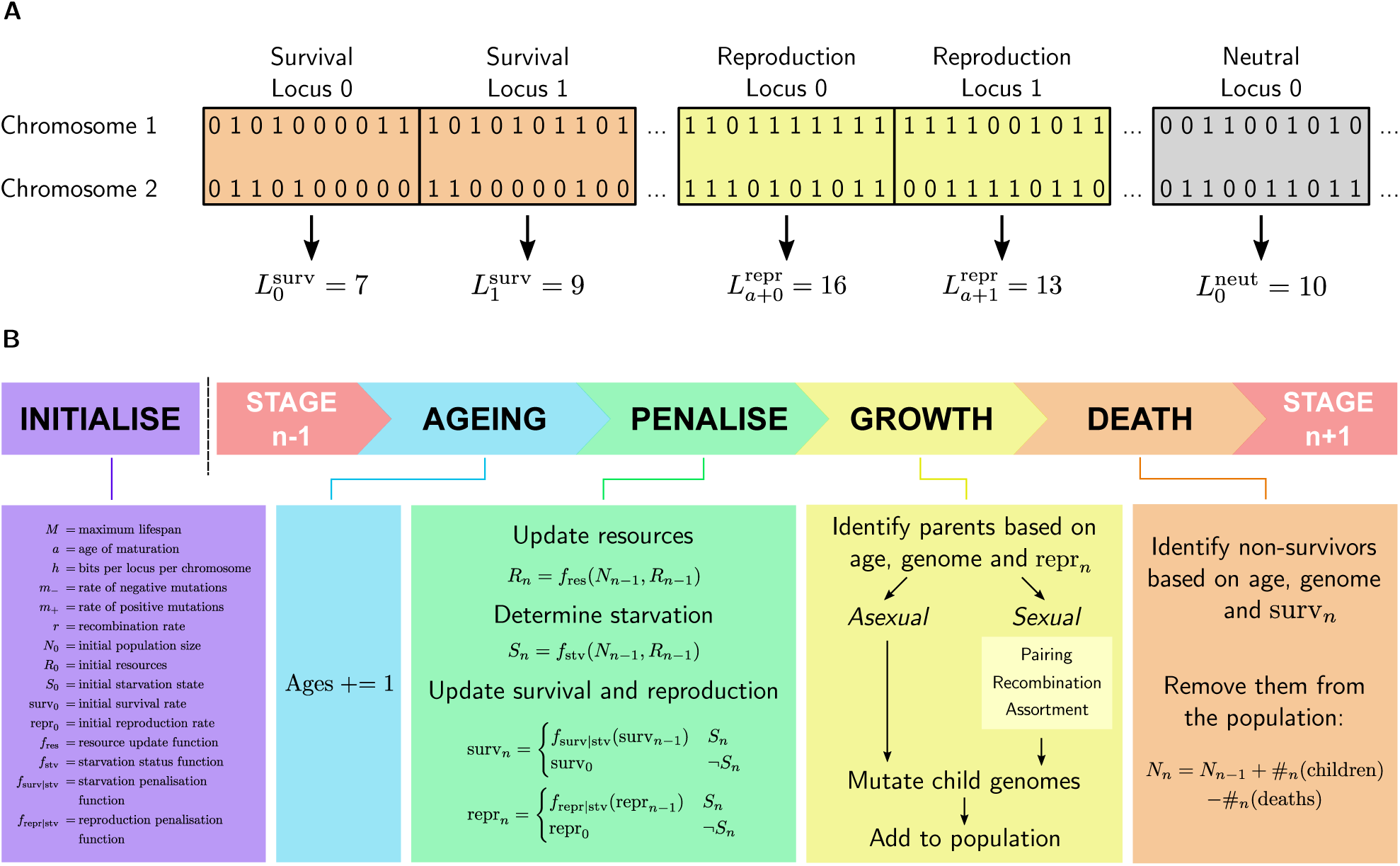
AEGIS workflow. (A) Each individual in an AEGIS population has a diploid bit-string genome comprising survival, reproduction and neutral loci. The sum *L* of bits across both chromosomes at a given locus position determines the probability of survival or reproduction at the appropriate age. *a* denotes the age of reproductive maturation for the population. (B) At each stage *n* of an AEGIS simulation, individuals increment their ages, then reproduce and die based on their ages, genomes and the starvation status of the population.

Upon initialisation, the population consists of some number of new individuals with uniformly-distributed genome composition and age values. The population is then permitted to evolve freely in discrete time, with individuals reproducing and dying at each stage according to the probabilities specified by their genomes (Fig. 1B). In asexual reproduction, each parent individual gives rise to one offspring per stage in which it reproduces, whose genome is first copied from the parent and then mutated. The rates of positive (0 *→* 1) and negative (1 *→* 0) mutations are specified separately; since mutations with a negative effect on fitness are much more common in real-world systems, the former probability is typically lower than the latter. In sexual populations, parent individuals are grouped randomly into pairs, the chromosomes of each parent undergo recombination with each other (Supplementary Material), and one chromosome is selected from each parent (assortment) to produce the child genome, which is mutated as above. In both cases, the allele composition of the new generation is drawn from that of the previous generation, and successive generations overlap within the population.

To limit the size of the population and impose competition between individuals, a resource limit is imposed on the population. By default, an initial resource level is set which remains constant throughout the simulation; if the size of the population exceeds this threshold, the survival probability of each individual is subjected to a compounding starvation penalty until the population falls below the resource limit. This typically leads to a rapid fluctuation of population size around the set value (Fig. 2A), as populations sequentially overshoot the resource limit and die back to a smaller size (Fig. 2B).

**Figure 2:**
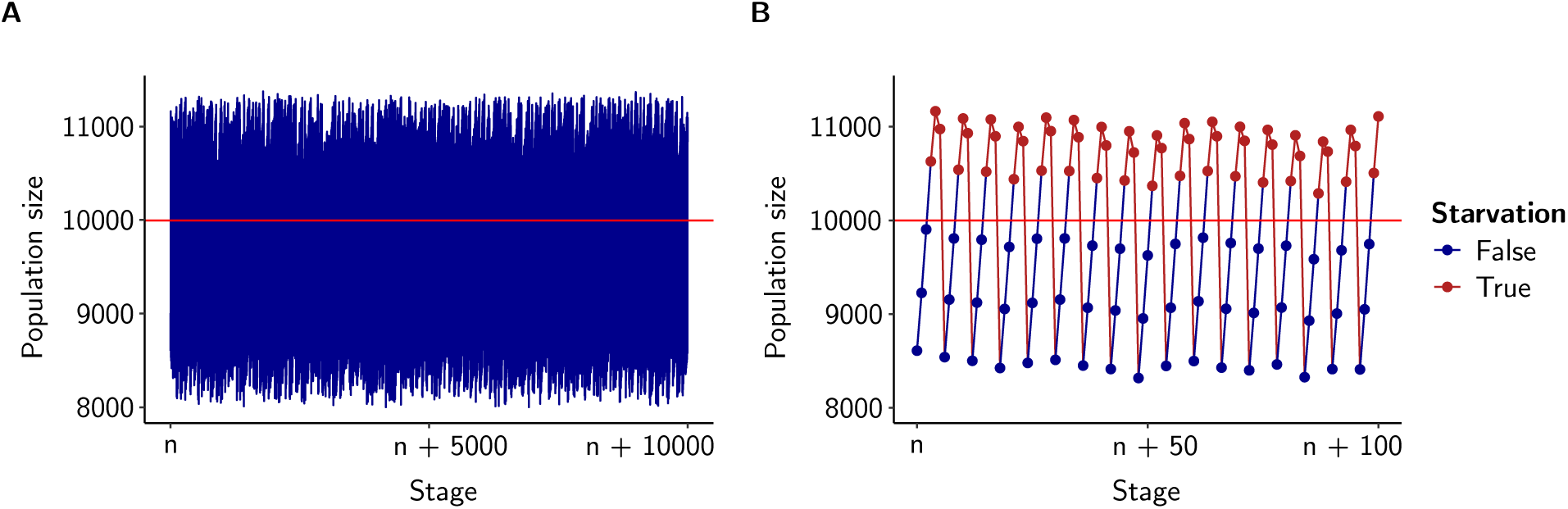
Population fluctuations in AEGIS simulations. (A) Trace of population size during 10,000 stages of an AEGIS run under sexual reproduction, showing cyclical fluctuations around a set resource level (horizontal red line), above which the population enters starvation. (B) Close-up trace of 100 stages from the same run, showing repeated cycles of population growth, starvation, and collapse.

One particularly important aspect of the model of evolution implemented by AEGIS is the manner in which it enables explicit calculation and comparison of fitness values. Because the baseline survival and reproduction probabilities of each individual are directly specified by its genome, the fitness of any individual (defined as its expected lifetime reproductive output) can be directly computed for any given set of probability bounds and starvation regime:

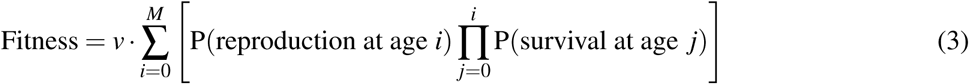

where *M* is the maximum lifespan of the population and *v* (equal to 1 for asexual populations and 0.5 for sexual ones) denotes the relative genetic contribution of a parent to its offspring. In the case of so-called *genotypic fitness*, this value is calculated directly using the baseline survival and reproduction probabilities specified by the individual’s genotype sums and user-specified probability bounds, without any starvation penalties. The distribution of individual and mean population fitness values can then be compared between populations to investigate the evolution of fitness in response to different conditions.

The runtime of an AEGIS simulation depends primarily on the number of stages, the population size (as determined by the resource limit), the genome size, and whether reproduction is sexual or asexual. Sexual reproduction is more computationally demanding, primarily due to the complexity of the recombination process. For example, a 256-GB-RAM machine with 8 CPUs per task was able to complete a one-million-stage simulation with the default genome size and asexual reproduction in 2 h 40 min when the resource limit was 1000 and 2 d 17 h 31 min when the resource limit was 20000; with sexual reproduction, the runtimes were roughly double this.

Following run completion, AEGIS can save a wide range of data in a cross-compatible CSV format. Some simple metrics, such as population size, are recorded at every stage of the run, while more complex information (such as genotype-frequency distributions) is recorded for a pre-specified number of “snapshot” stages evenly distributed throughout the run. The data output by AEGIS can be used for a wide variety of downstream analysis and visualisation purposes.

A detailed tutorial for AEGIS installation and use is provided along with example configuration files at github.com/valenzano-lab/aegis.

## Ageing evolves differently in sexual and asexual populations

One of the most fundamental applications of the AEGIS simulation tool is in investigating the evolution of age-dependent survival and reproduction across different conditions. Starting from an initial genome containing uniformly-distributed 0’s and 1’s, we find that ageing-like phenotypes reliably evolve across a wide range of population sizes, mutation rates and reproductive strategies (Fig. 3A to 3C). When mutation rates are sufficiently low, loci determining survival and reproduction in early life accumulate large numbers of beneficial mutations, resulting in low baseline (i.e. non-starvation) mortality rates before and immediately after reproductive maturation and high fecundity levels in early adulthood. Following reproductive maturation, survival and reproduction rates progressively decline as the genotype sums of the corresponding loci accumulate progressively larger numbers of deleterious mutations. Remarkably, the genotype sums of loci affecting older age groups consistently converge on the mean genotype value of the neutral loci in the genome, indicating that selection is relaxed towards neutrality in genes affecting late-life.

**Figure 3:**
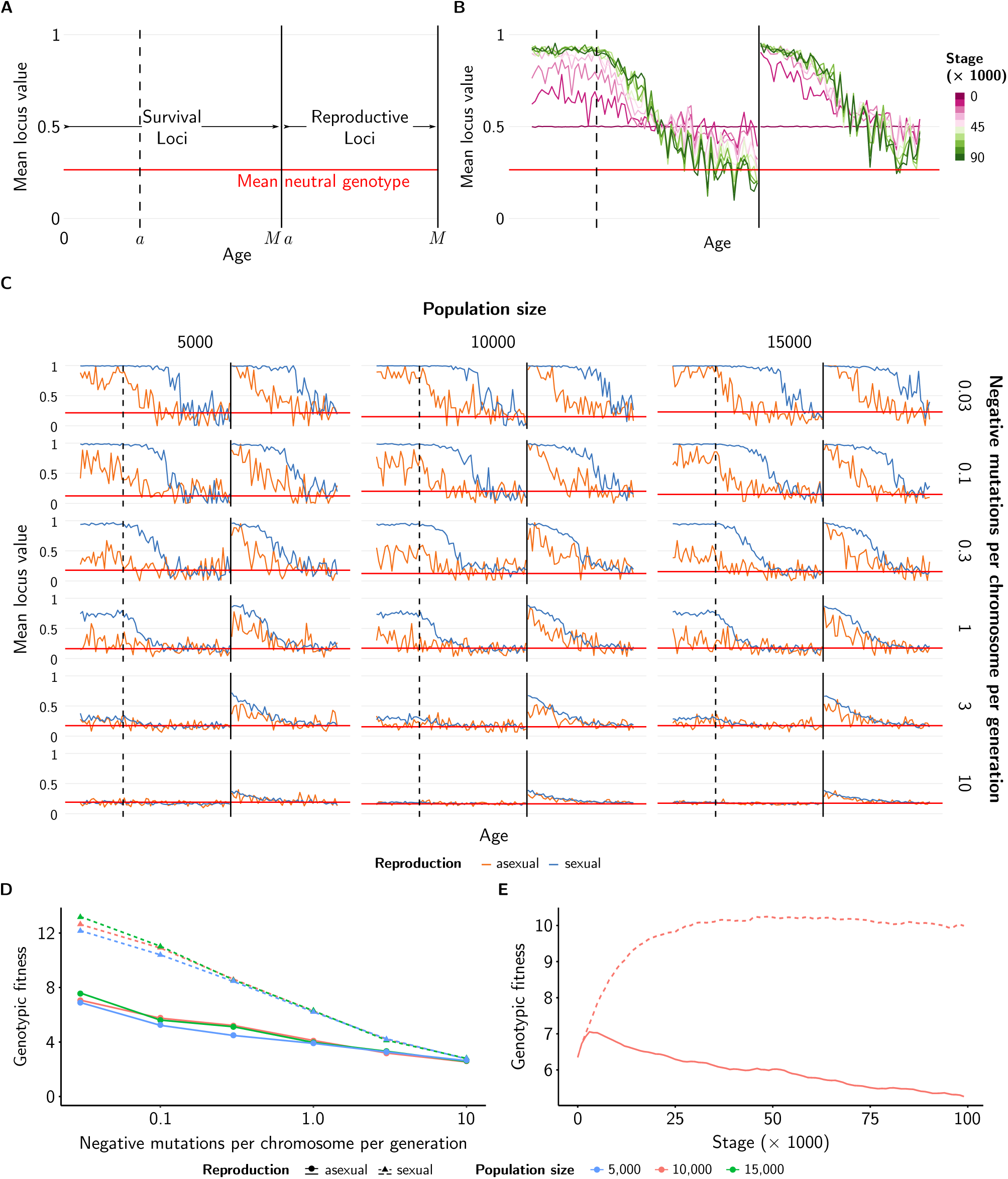
Life-history evolution in the AEGIS model. (A) Explanation of genotype plots in subsequent panels. Loci coding for survival from age 0 to maximum lifespan, *M*, are plotted in order on the left, while loci coding for reproduction probabilities from reproductive maturation, *a*, to maximum lifespan are shown on the right of the solid vertical line. The dashed vertical line indicates the transition from pre-maturation to post-maturation survival loci. The red horizontal line shows the mean value of neutral loci across all populations shown on a given pair of axes. (B) Genotype plot of a sexual population under a negative mutation rate of 0.3 per chromosome per generation and a resource level of 10,000, showing the progressive evolution of an ageing phenotype over 100,000 stages. (C) Grid of genotype plots showing the state of sexual and asexual populations under a variety of resource and mutation conditions after 1 million stages. (D) Plot of average genotypic fitness values for the same populations after 1 million stages, showing the decline in genotypic fitness with increasing mutation rate. (E) Plot of average genotypic fitness of a sexual and an asexual population under a negative mutation rate of 0.3 and a resource level of 10,000, over the first 100,000 stages of a simulation. In all subfigures, *a* = 21, *M* = 70, and the rate of positive mutations is equal to 20% of the rate of negative mutations.

While ageing-like phenotypes consistently evolved across a wide range of initial conditions, the specific outcome of the simulation depended heavily on the mutation rate and reproductive strategy imposed on the population (Fig. 3C). At very low mutation rates, pre-reproductive-maturation survival rates evolve to near-maximal levels, and very high baseline survival and reproduction probabilities often persist for extended periods following maturation. As mutation levels increase, the pre-maturation survival rates and the post-maturation decline in survival and reproduction shift to progressively earlier ages. At very high mutation rates, the increased survival and reproduction of early ages is completely abrogated, and the entire genome behaves similarly to the neutral loci. Hence, as the mutation rate increases, the level of selection required to maintain a favourable genotype increases, resulting in a shift towards more rapid ageing and shorter expected lifespans.

In addition to the effect of mutation rates, the choice between sexual and asexual reproduction has dramatic effects on the evolution of ageing in the AEGIS model. Under the rates of mutation and recombination used in Fig. 3, asexual populations consistently exhibit lower pre-maturation survival rates and more rapid post-maturation declines in survival and reproduction rates than sexual populations (Fig. 3C). As a result, survival and reproduction rates in sexual populations typically exceed those of asexual populations evolving under similar conditions, and the transition from a condition of elevated early-life fitness to one in which the entire genome appears to evolve neutrally occurs at lower mutation rates when reproduction is asexual.

As a result of these differences in life history evolution, the average genotypic fitness of individuals in sexual populations consistently evolves to a higher level than in asexual populations, with the size of the gap increasing as the mutation rate declines (Fig. 3D); only at very high mutation rates, at which both reproductive strategies give rise to near-neutral reproduction and survival phenotypes at most loci, do the fitnesses of sexual and asexual populations converge. Investigating genotypic fitness at different points in time under intermediate mutation rates (Fig. 3E) reveals the kinetics of this divergence: although both sexual and asexual populations begin at the same average genotypic fitness, in sexual populations the genotypic fitness progressively increases to a high equilibrium value, while in asexual populations a small initial increase is followed by progressive decay, in a manner compatible with the accumulation of irreversible mutations predicted by Muller’s ratchet (Muller, 1964; Felsenstein, 1974).

Why would sexual and asexual populations evolve such different life histories under shared environmental, genetic and phenotypic constraints? One plausible explanation is the Hill-Robertson effect (Hill et al., 1966), whereby recombination and assortment enable beneficial mutations occurring in different lineages to accumulate on the same chromosome. In contrast, each asexual individual is restricted to mutations occurring within its single line of ancestors, and improvements to population fitness can only occur through competition between autarkic asexual lineages. As a result, sexual populations are able to accumulate larger numbers of beneficial mutations in loci affecting early-life survival and reproduction relatively rapidly, and can sustain higher rates of survival and reproduction in the face of a given rate of negative mutations. Under this explanation, the differences between life histories evolved by sexual and asexual populations are therefore driven primarily by differences in positive, rather than purifying, selection.

## Discussion

The evolutionary mechanisms underlying the widespread occurrence of senescence across taxa have long been a topic of interest among evolutionary biologists, population geneticists, and biogerontologists. Genomic surveys in unusually long- or short-lived species have attempted to identify the genetic changes underlying differences in life histories across species, typically by identifying genes exhibiting significant sequence changes in particular taxa (Keane et al., 2015; Kim et al., 2011; Seim et al., 2013; Valenzano et al., 2015) potentially associated with positive selection. While experimental work of this kind has identified specific genes and conserved molecular pathways impacting ageing and lifespan in particular species (Tacutu et al., 2017), little is known about how differences in life history evolve between natural populations. To date, except for a few simple models (Stauffer, 2007), there has been a general lack of numerical tools for simulating the evolution of life histories, impeding the investigation of how ageing and lifespan evolve under different selective conditions.

AEGIS is intended to fill this gap, providing a versatile and accessible numerical tool to simulate the evolution of lifespan and ageing under a wide range of genetic, selective and demographic constraints. The AEGIS software is simple to install and use, can run on both personal computers (for simple runs) and high-performance clusters (for large, intensive runs) and provides ready-to-use graphical visualisations and cross-platform output for downstream investigations. Since survival and reproduction probabilities are explicitly encoded in the genomes of AEGIS populations, the model allows accurate calculation of individual and mean population fitness, enabling the investigation of time-dependent changes in allele frequency under different selective constraints.

While AEGIS exceeds previous models in its flexibility and power, there are nevertheless a number of important extentions and improvements that could be made. The current AEGIS model contains no scope for inter-population competition, freely-evolving mutation rates, or the multi-age-affecting loci that would be needed to test theories of ageing relying on epistatic or pleiotropic effects. Future work on the model, both by the current authors and other contributors to the open-source AEGIS project, will fill these gaps and further improve our ability to use numerical simulation to interrogate the evolution of ageing.

## Acknowledgements

We are thankful to all the members of the Valenzano lab at the Max Planck Institute for Biology of Ageing for continuous feedback on this project and to Jorge Boucas and all the members of the MPI-Age bioinformatic core facility and IT for allowing continuous usage of the MPI-Age computing cluster. We would like to thank Fabio Iocco for discussions and intellectual contribution to the early stages of development of this project. This work has been entirely supported by the Max Planck Society and the Max Planck Institute for Biology of Ageing in Cologne, Germany.

## Supplementary Material

### The AEGIS run

#### Initialisation

Every AEGIS simulation is initialised from a configuration file, which specifies the population and run parameters for that simulation (the config file for Fig. 3B, for example, is shown in Fig. S1). Upon run initialisation, the parameters in the config file are used to derive a range of other parameters, such as the survival and reproduction probabilities corresponding to each possible genotype, the number of loci, and the length of each chromosome in bits. The per-bit probability *m*_+_ of positive mutations is determined based on the user-specified probability *m*_−_ of negative mutations and the specified positive:negative mutation ratio *v*:

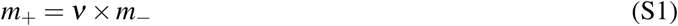

A single genome layout is defined for all individuals, and is randomised at the start of the simulation to misimise the impact of interlocus linkage effects. Finally, the starting population of individuals is initialised: by default, the starting genomes of the population are drawn from a discrete uniform distribution with sample space {0, 1}, while the starting age of each individual is sampled randomly from the set {0, 1, …, *M*}, where *M* is the maximum lifespan.

In the case of the simulations presented in Fig. 2 and 3, the user-defined bounds on survival and reproduction probability (from which age-dependent probabilities are derived via Equations 1 and 2) were defined such that firstly, the population does not regularly go extinct over the course of the simulation, and secondly, almost all individuals die out before reaching maximum lifespan.

#### Stage progression

Following initialisation, the run progresses in discrete time stages. At the start of each stage, the age of each individual increments by 1. The available resources are then updated, and the starvation state of the population is determined based on the population size and the available resource level: by default, resources are constant and the population enters starvation if the population size exceeds the set resource level. If starvation occurs, the survival and reproduction probabilities corresponding to each genotype sum value are penalised based on the number of turns the population has been in starvation; by default the probability of death trebles each turn during starvation, while reproduction probability remains constant. The imposition of a starvation penalty in this manner limits the size of the population, and thus ensures the simulation remains practically computable, while also imposing competition for resources between individuals.

Following resource updating, the population enters the reproduction phase. For each individual, the probability of parental status is determined based on its age, the genotype sum of the reproduction locus corresponding to that age, and the user-specified probability bounds (Equation 2). Each parent is then randomly and independently classified as a parent or non-parent based on this probability. In asexual populations, the population of parents is then duplicated to generate a population of children, each of which is assigned an age of 0. The genome of each child is then mutated according to the probabilities of positive and negative mutations determined during initialisation: each 0-bit is independently mutated to a 1-bit with probability *m*_+_, and each 1-bit is independently mutated to a 0-bit with probability *m*_−_. Finally, the population of children is then added to the overall population.

In sexual populations, the population of parents is grouped into mating pairs at random; in the event of an odd number of parents, one parent is selected at random and does not reproduce. The two chromosomes of each parent undergo recombination (see below), shuffling corresponding genome segments between the two chromosomes. After recombination, one chromosome is selected at random from each parent, and the two chromosomes are concatenated in a random order to generate a new child individual (assortment). Each child produced in this way is assigned age 0, and the genome of the child population is mutated as in the asexual case before being added to the overall population.

**Figure S1:**
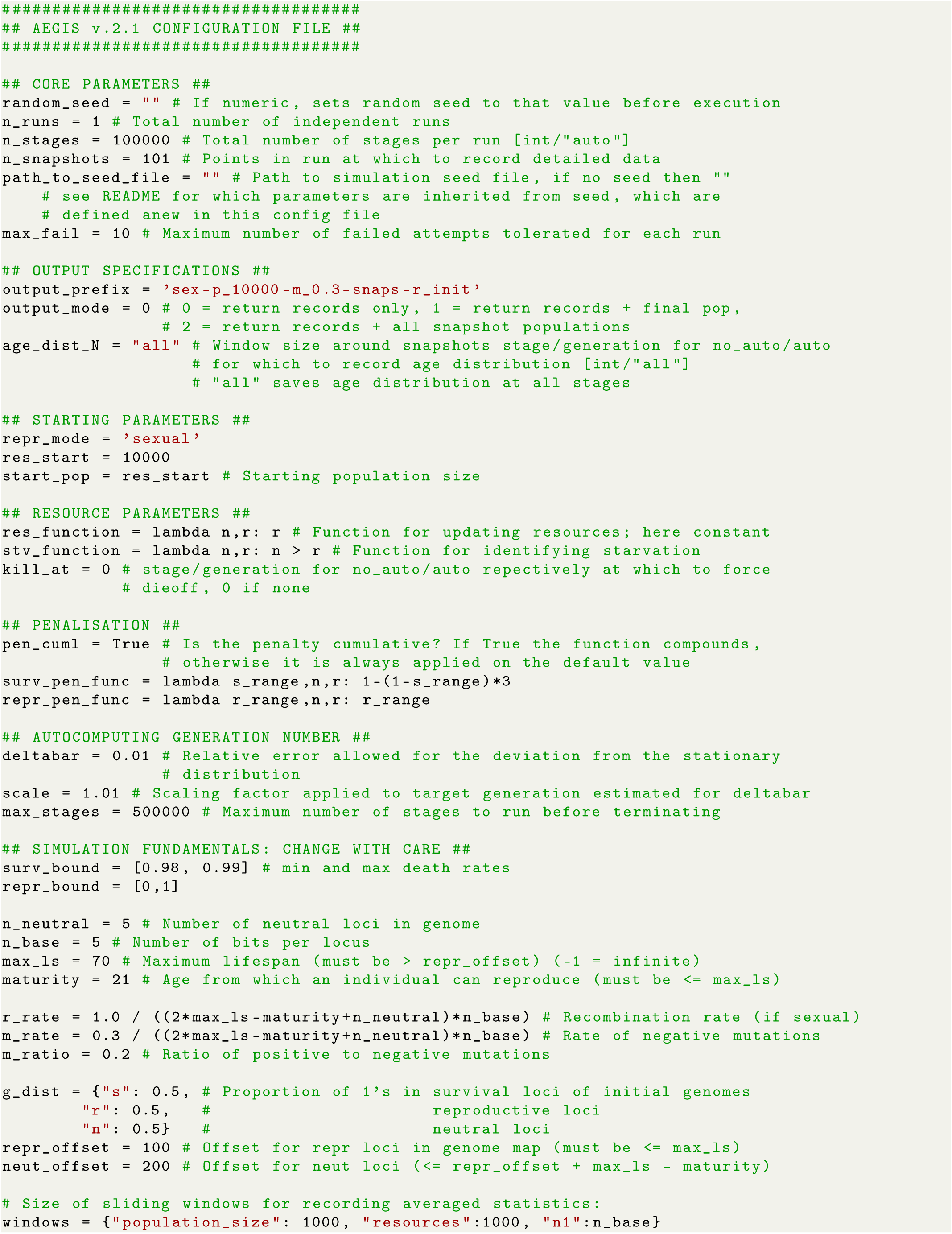
An example AEGIS config file.

Following reproduction, the final phase of each stage is death of individuals. As in reproduction, the survival probability of each individual is determined based on its age, the genotype sum of the corresponding locus, and the user-specified probability bounds (Equation 1), and each individual is independently classified as surviving or dying based on these probabilities. The population of survivors is retained for the next stage of the simulation, while the remaining individuals are discarded.

The complete code of the AEGIS simulation software, including the functions for all the above operations, is available at github.com/valenzano-lab/aegis.

#### Recombination

Conceptually, recombination involves aligning the homologous chromosomes of an individual and randomly exchanging corresponding portions of sequence between each chromosome pair. Computationally, this process can be simulated by randomly determining recombination sites along the length of the chromosome (with some independent probability *r* for each position to be selected as a recombination site) and exchanging the corresponding sequence on each chromosome between each site and the end of the chromosome (Fig. S2A). However, rather than actually performing a large number of exchange operations, it is computationally far more efficient to simply count the number of recombinations affecting each position along the chromosome and exchange only those portions affected by an odd number of recombination events (Fig. S2B).

In order to avoid a directional bias in recombination (in which, for example, positions at the end of the chromosome are much more likely to be transferred between chromosomes than positions at the beginning), it is also important to randomly determine the orientation of each recombination event, such that some events affect sequence between the recombination site and the end of the chromosome and others affect sequence between the start of the chromosome and the recombination site (Fig. S2C).

In AEGIS, therefore, recombination in sexual populations is implemented as two independent processes, one producing forward-oriented recombination events and the other reverse-oriented ones. At the beginning of the recombination process, forward and reverse recombination sites are determined randomly, with each site having an independent 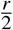 probability of being selected as a forward site and the same probability of being selected as a reverse site. Each event affects all chromosome positions between the recombination site and the appropriate end of the chromosome, inclusive of the recombination site. The number of recombination events affecting each chromosomal position is then quantified, and regions of sequence affected by an odd number of events are exchanged between the two chromosomes (Fig. S2B and S2C).

At present, interference between recombination sites is not implemented in AEGIS. Modification of the recombination algorithm to permit different interference functions could be implemented in a future extension of the software.

#### Analytic behaviour of the neutral loci

A locus in the AEGIS genome without a phenotypic (i.e. reproduction or survival) effect is referred to as *neutral*. As these loci are not affected by selection on survival or reproduction rates, their evolution over time is relatively easy to model analytically, especially in the asexual case, and provides some useful predictions about the behaviour of the model over time.

**Figure S2:**
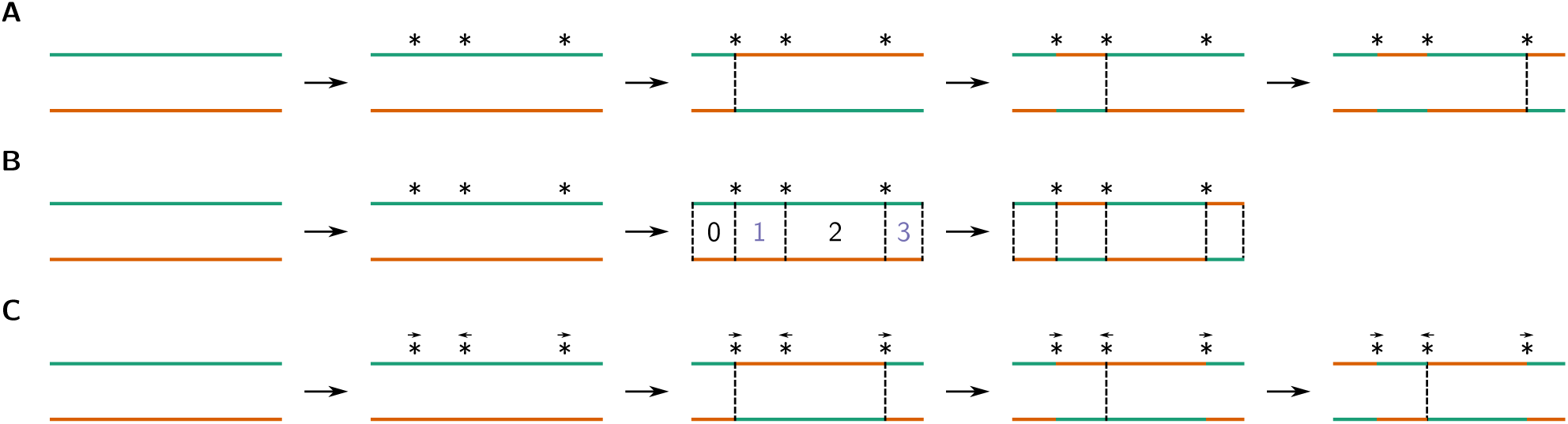
Recombination in the AEGIS model. (A) In a naïve implementation of chromosomal recombination, recombination sites are determined at random and corresponding chromosomal sequences are exchanged at each site in turn. (B) In a more efficient implementation, the number of recombination events affecting each chromosomal position is counted, and regions affected by an odd number of events are exchanged. (C) In order to avoid directional bias in the probability of a given chromosomal position being exchanged between chromosomes, forward- and reverse-oriented recombination events must occur with equal probability.

Unlike with survival and reproduction loci, the *sum* over the bits in a neutral locus has no phenotypic effect; as a result, in the absence of linkage, the evolution of each bit in the locus can be assumed to evolve independently.

Let *µ* denote the rate of negative (1 *→* 0) mutations and *v* the ratio of positive to negative mutations. The rate of positive (0 *→* 1) mutations is then given by *µ · v*. These transition probabilities are memoryless: conditional on the state of the bit, the probability of a transition is independent of its past states. The evolution of each bit in the neutral locus can therefore be modelled as a two-state discrete-time Markov chain, with transition matrix

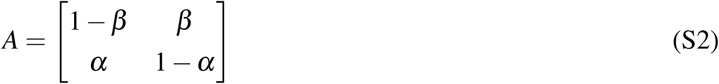

where *α* = *µ, β* = *µv*, the first row and column indicate state 0 and the second row and column indicate state 1. At generation *k*, the state distribution of the Markov chain is therefore given by

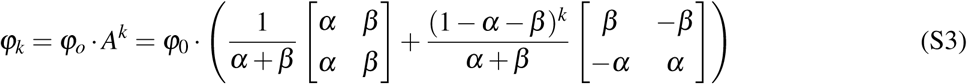

After many generations of mutation, the second term within the parentheses tends towards zero, and the Markov chain approaches its limiting distribution:

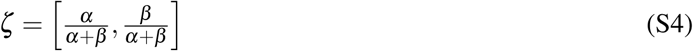

As a result, the expected value of the bits in a neutral locus converges over time to

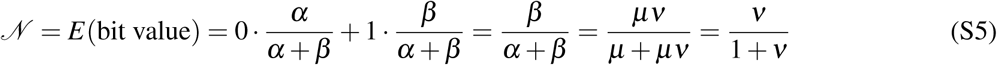

The equilibrium mean value of neutral loci, therefore, is independent of the mutation rate, and depends only on the ratio between positive and negative mutations. When *v* = 0.2, for example, *𝒩* converges to a value of 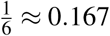, as can be observed in the genotype plots in Fig. 3C.

We can go further and calculate the degree to which the average value *p*_*k*_ of the neutral locus at any generation *k* deviates from this equilibrium value *𝒩*:

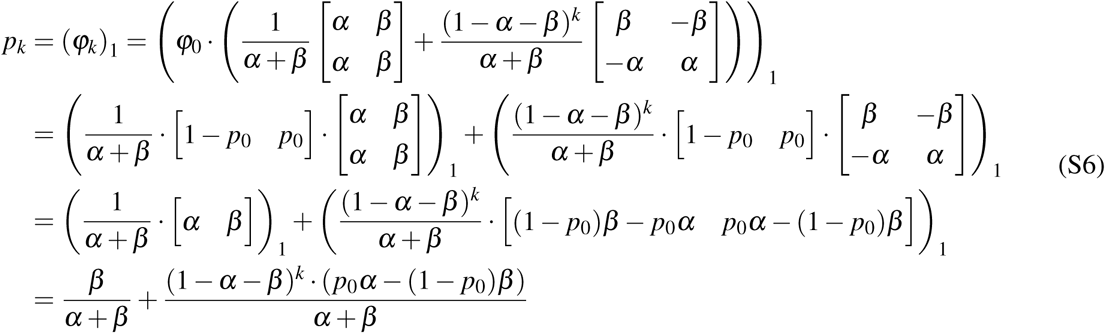

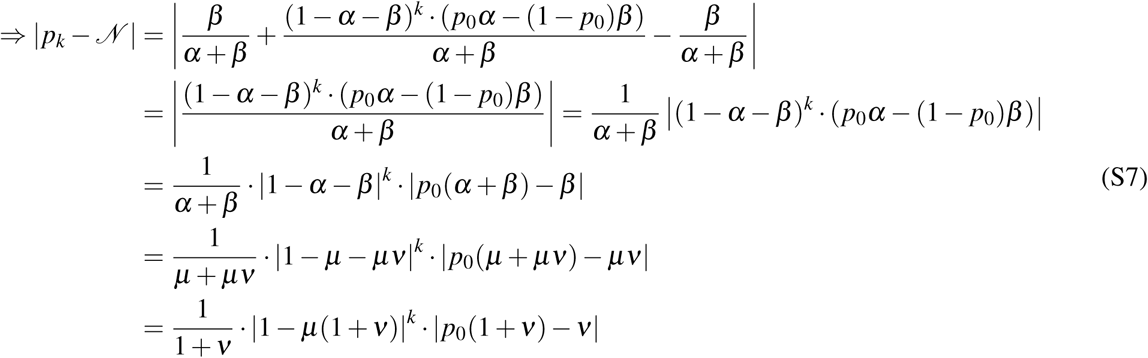

For any deviation *δ* between *p*_0_ and *𝒩,* therefore, we can calculate the earliest generation *K* for which *|p*_*k*_ − *𝒩 | < δ*:

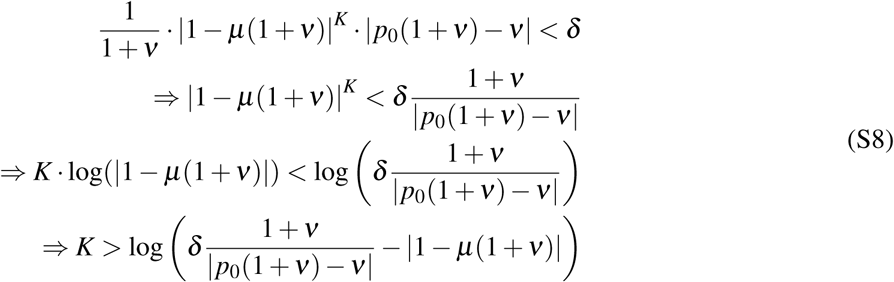

Unlike the value of *𝒩* itself, therefore, the number of generations required for *p*_*k*_ to converge to within a given distance of *𝒩* does depend on the mutation rate *µ*, with higher values of *µ* resulting in faster convergence times.

The above calculations provide an alternative method for specifying the number of stages in an AEGIS run: rather than explicitly specifying a total number of stages, one can specify the desired value of *δ,* and the simulation will run until all individuals in the population are at least *K* generations removed from the starting population, then stop. In principle, this provides a more reliable method for ensuring that the population has evolved to a sufficient state of equilibrium; however, for simplicity, and due to the extra complications introduced by the sexual case (which are not covered here), we have restricted ourselves in this publication to simulations running for a fixed number of stages.

